# Identification of a global gene expression signature associated with the genetic risk of catastrophic fracture in iPSC-derived osteoblasts from Thoroughbred horses

**DOI:** 10.1101/2024.06.19.599695

**Authors:** Esther Palomino Lago, Amy K. C. Ross, Alyce McClellan, Deborah J. Guest

## Abstract

Bone fractures are a significant problem in Thoroughbred racehorses. The risk of fracture is influenced by both genetic and environmental factors. To determine the biological processes that are affected in genetically susceptible horses, we utilised polygenic risk scoring to establish induced pluripotent stem cells (iPSCs) from horses at high and low genetic risk. RNA-sequencing on iPSC-derived osteoblasts revealed 112 genes that were significantly differentially expressed. 43 of these genes have known roles in bone, 27 are not yet annotated in the equine genome and 42 currently have no described role in bone. However, many of the proteins encoded by the known and unknown genes have reported interactions. Functional enrichment analyses revealed that the differentially expressed genes were overrepresented in processes regulating the extracellular matrix and pathways known to be involved in bone remodelling and bone diseases. Gene set enrichment analysis also detected numerous biological processes and pathways involved in glycolysis with the associated genes having a higher expression in the iPSC-osteoblasts from horses with low polygenic risk scores for fracture.

Therefore, the differentially expressed genes may be relevant for maintaining bone homeostasis and contribute to fracture risk. A deeper understanding of the consequences of mis-regulation of these genes and the identification of the DNA variants which underpin their differential expression may reveal more about the molecular mechanisms which are involved in equine bone health and fracture risk.

## Introduction

Bone fractures in horses can occur due to traumatic events (such as a kick or fall), but in Thoroughbred racehorses, they often occur in the absence of any trauma and in response to bone overloading. Fractures due to bone overloading are the most common musculoskeletal injury in training and racing accounting for 41% of all musculoskeletal injuries (Johnston et al., 2020a). Non-fatal stress fractures lead to a significant loss of time in training, earnings and race starts (Johnston et al., 2020b). However, more complex fractures in horses can be difficult to treat because of the need for continuous weight-bearing in the injured limb during recovery. As such, fracture is the main reason for euthanasia on the racecourse (McKee, 1995), with an average of 60 horses/year suffering a fatal, catastrophic distal limb fracture during racing in the UK (Parkin et al., 2004a). Fractures therefore have a large economical and welfare impact.

Many environmental risk factors for fracture have been identified (Georgopoulos and Parkin, 2017, Parkin et al., 2004b, Verheyen et al., 2006, Anthenill et al., 2007, Kristoffersen et al., 2010) but a genetic component to fracture risk has also been shown. The heritability of fracture risk in Thoroughbreds is 0.21-0.37 (Welsh et al., 2014). A genome wide association study (GWAS) identified significant genetic variation associated with catastrophic fracture risk on chromosomes 9, 18, 22 and 31 and four significantly associated SNPs; three on ECA18 (horse chromosome 18) and one on ECA1 (Blott et al., 2014). Additional SNPs within the region on ECA18 were also found to be associated with fracture (Tozaki et al., 2020). Studies on the pathology of fracture indicate stress-related damage to the bone prior to catastrophic injury (Stover, 2003). This may be related to abnormal bone repair, as horses euthanised following a fracture show changes in their other bones including stress fractures and excessive bone remodelling (Anthenill et al., 2010, Parkin et al., 2006, Stover et al., 1992, Riggs et al., 1999). This suggests that the bone cells involved in tissue homeostasis may be affected in horses that suffer from fatal fractures. However, it has proven challenging to understand the genes and biological pathways that may be involved in fracture risk and studies using bone tissues are complicated by the confounding environmental risk factors.

We have recently developed a genome-wide polygenic risk score (PRS) for catastrophic fracture (Palomino Lago et al., 2024). PRS provide an estimate of the genetic risk of the phenotypic trait at the level of the individual (Choi et al., 2020) and their potential clinical utility is becoming increasingly evident (Lewis and Vassos, 2020). We previously utilised the PRS for fracture to establish an *in vitro* cell model to study bone gene regulation in cells from horses at high and low risk (Palomino Lago et al., 2024), in the absence of any environmental variability. In that study we used a candidate gene approach and identified that *COL3A1* was differentially expressed between high and low risk horses, in part due to the presence of an upstream SNP that was significantly associated with fracture.

However, the candidate gene approach only provided insights into a very narrow range of genes and the use of primary cells in this model was associated with technical limitations such as restricted cell growth. Induced pluripotent stem cells (iPSCs) may provide better cell models due to their ability to differentiate into a wide range of cell types and unlimited growth *in vitro*. They have been used to study many inherited conditions in humans (Mae et al., 2023, Fear et al., 2023, Gorashi et al., 2023) and are of particular value when access to the affected cell types is limited and/or the cells do not expand well in culture (Pereira et al., 2023). More recently human iPSCs have been used in conjunction with polygenic risk scores (Dobrindt et al., 2021, Coleman, 2022, Yde Ohki et al., 2023) to study the mechanisms underlying complex genetic conditions. We have previously derived iPSCs from horses (Bavin et al., 2015, Palomino Lago et al., 2023) and established the methods to differentiate them into bone forming osteoblasts (Baird et al., 2018, Baird et al., 2019). Furthermore, we demonstrated that the genetic background of the iPSCs affects their expression of osteoblast-associated genes, whereas different iPSC-lines derived from the same horse had less variance (Baird et al., 2018).

The aim of this study was to use iPSC-osteoblasts derived from Thoroughbred horses with high and low PRS for catastrophic fracture and perform global gene expression analysis to identify the biological processes and pathways that are perturbed in osteoblasts from high risk horses.

## Material and Methods

### Experimental design

An overview of the experimental design used in this study is shown in Fig. 1. Frozen stocks of skin fibroblasts that had been derived and banked from six male Thoroughbred horses were used. The skin biopsies were taken at post-mortem from horses which had been euthanised for reasons unrelated to this study and with the consent of the Animal Health Trust Ethical Review Committee (AHT_02_2012) and Royal Veterinary College Clinical Research Ethical Review Board (URN 2021 2035-2). The samples had previously been genotyped and polygenic risk scores for catastrophic fracture had been calculated to select samples representing the high and low ends of the risk spectrum (Palomino Lago et al., 2024).

**Fig 1.**
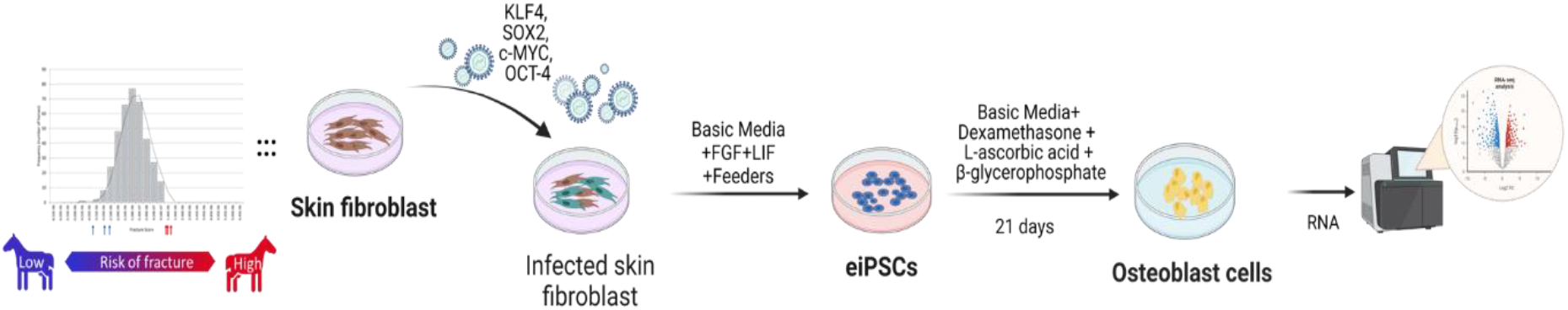
Overview of the experimental approach. Created with Biorender.com.

### iPSC generation

Induced pluripotent stem cells (iPSCs) lines were generated previously from skin fibroblasts by retroviral transduction using methods previously described (Bavin et al., 2015, Baird et al., 2018). Briefly, phoenix gag-pol packaging cells were transfected with 3 µg of pVPack-VSV-G (Agilent technologies, UK) along with 3 µg of pMXs.hOct4 (Octamer-Binding Protein 4, RRID:Addgene_17217), pMXs.hSox2 (SRY-Box Transcription Factor 2, RRID:Addgene_17218), pMXs.hKlf4 (Kruppel Like Factor 4RRID:Addgene_17219), pMXs.hc-Myc (MYC Proto-Oncogene, RRID:Addgene_17220), or pMX.GFP (Cell Biolabs, USA). Transfections were carried out using lipofectamine3000 and Opti-MEM media (both Invitrogen, Thermo Fisher, UK) according to the manufacturer’s instructions. After 48 h culture supernatant containing the viral particles was pooled, filtered through a 0.45 µM filter (Nalgene, Thermo Fisher, UK), supplemented with 10 µg/ml polybrene (Sigma-Aldrich, UK) and used to infect equine skin fibroblasts which had been plated at a density of 1×10^4^ the day before infection. After three rounds of viral infection, at 48 h intervals, infected cells were plated at a density of 5×10^3^ cells per 10 cm plate pre-seeded with feeder cells (mitotically inactivated mouse embryonic fibroblasts). The media was replaced with basic media [DMEM/F12, 15% fetal bovine serum (FBS), 2 mM L-glutamine, 0.1 mM non-essential amino acids, 0.1 mM 2-βmercaptoethanol, 1 mM sodium pyruvate (all from Thermo Fisher, UK)] plus 1000 U/ml Leukemia inhibitor factor (LIF) (Preprotech, UK) and 10 ng/ml basic fibroblast growth factor (bFGF) (Peprotech). iPSC media was replaced every other day until iPSC colonies began to appear and reached a large enough size to manually pick selected colonies. These colonies were used to establish clonal iPSC lines. For each skin fibroblast sample, two or three clonal lines of iPSCs were generated to provide a total of 14 iPSC lines. iPSCs were mechanically passaged in the presence of 2 μM Thiazovivin (Miltenyi Biotec, UK).

### iPSC differentiation into osteoblasts

Osteoblasts were generated as previously described (Baird et al., 2018). Small pieces of iPSC colonies were plated at a density of 7×10^4^ cells per well of a 24-well OsteoAssay surface coated plate (Corning, Wiesbaden, Germany) without feeders and in iPSC media in the presence of 2 μM Thiazovivin for the first 24 h. The following day, the media was replaced with osteogenic media [DMEM/F12, supplemented with: 15% FBS, 2 mM L-glutamine, 1% non-essential amino acids, 1 mM sodium pyruvate, 0.1 mM 2-mercaptoethanol (all Invitrogen, Thermo Fisher, UK), 10 mM β-glycerophosphate, 10 nM dexamethasone and 28 µM ascorbic acid (all Sigma-Aldrich)]. Cells were cultured for 21 days with the media replaced every 2–3 days prior to RNA extraction.

### RNA extraction

Following cell differentiation, RNA was collected using Tri-reagent (Sigma-Aldrich) and extracted using the RNeasy Mini Kit (Qiagen, UK) following the manufacturer’s instructions. Purified RNA was treated with the DNA-free™ DNA removal kit (Invitrogen, Thermo Fisher) to remove genomic DNA contamination according to the manufacturer’s instructions. RNA concentration was determined using a Qubit (Thermo Fisher Scientific) and the purity was determined using a DS-11 spectrophotometer to ensure 260/280 ratios >1.8. RNA integrity was measured using a Tapestation (2100 Bioanalyzer (Agilent, UK) and for all samples was confirmed to be >9.0.

### RNA sequencing

RNA from 14 lines of iPSC-osteoblasts (passage 8-15) was used in RNA sequencing. mRNA library preparation and transcriptome sequencing were conducted by Novogene (Cambridge, UK) using an Illumina NovaSeq 6000 to generate 26.1-57.2 million 150 bp paired end reads per sample. FastQC (v.0.11.9) (Babraham Bioinformatics, Cambridge, UK) and MultiQC (v1.11) were used to establish raw sequencing quality. Raw RNA-seq reads were aligned to equine transcriptome obtained from NCBI EquCab 3.0 (GCF_002863925.1_EquCab3.0) annotation release 103 using the pseudoaligner Salmon (v.1.5.2) in Quasi-mapping-based mode with GC-bias correction (Patro et al., 2017). Then, transcript level mappings were imported into RStudio (v.4.2.1) using Tximport (v.1.20) (Soneson et al., 2015). Finally, read counts were normalized and analysed for differential expression using DESeq2 v.1.22.2 (Love et al., 2014). Genes with a Log2Fold Change (Log2FC) of ±1 and an adjusted p-value (p-adj) of ≤0.05 were considered as differentially expressed. P-values were adjusted using the Benjamini and Hochberg method in DESeq2.

### Functional annotation analysis

To investigate the biological function related to the DEGs, DAVID v.6.9 (Huang et al., 2009) and the PANTHER classification system (https://www.pantherdb.org/) v.17.0 were used to functionally annotate genes based on Gene Ontology (GO) terms (BP, biological process, MF, molecular function, CC, cellular component) and pathways. False Discovery Rate (FDR) < 0.05 was considered to be a statistically significant enrichment. Gene Analytics from the LifeMap’s GeneCards Suite (https://geneanalytics.genecards.org/) was used to perform pathway analysis, with an entity score of >6 being equivalent to a corrected p-value of ≤0.05 and therefore defined as significantly enriched. The results of the GO and Pathway analyses were visualized using R (v.4.2.2).

### Protein-protein interaction (PPI) network construction

Network analysis was conducted using the STRING (v.10.5) protein network analyser plug-in of Cytoescape (v3.9.1) (Shannon et al., 2003).

### Gene set enrichment analysis (GSEA)

Functional Class Sorting (FCS) of all expressed genes, regardless of whether they were significantly differentially expressed, was performed using Gene Set Enrichment Analysis (GSEA) software (v.4.3.2) based on H (Hallmark gene sets), C2 (curated gene sets): REACTOME, C2:WP, C2:KEGG, C5 (ontology gene sets) gene set collections (MSigDB v.2023.1) (Subramanian et al., 2005). GSEA first ranked all expressed genes according to the significance of differential gene expression between the high risk and low risk of fracture groups. The enrichment score for each gene set is then calculated using the entire ranked list, which reflects how the genes for each set are distributed in the ranked list. Normalized enriched score (NES) were determined for each gene set. The significant enrichment of gene set cut-off was |NES| >1, nominal p-value ≤ 0.01, and FDR<0.25.

### cDNA synthesis and quantitative RT-PCR

To validate the RNAseq, qPCR was used to measure a number of genes using the same samples as had been used in the RNAseq. cDNA was synthesised from 1 µg of RNA using the sensiFAST cDNA synthesis kit (Bioline, UK). 2 µl of cDNA (corresponding to 20 ng) was used in qPCR. Equine specific primers were designed using PrimerBlast (https://www.ncbi.nlm.nih.gov/tools/primer-blast/index.cgi) and mfold (http://www.unafold.org/) to produce amplicons of 50-150 bp. Primer sequences can be found in Supplementary Table S1. Quantitative PCR (qPCR) was performed in duplicate using SYBR Green containing supermix (Bioline) on a Bio-rad C1000 Touch Thermal Cycler (Bio-rad, UK). PCR cycle parameters were as follows: 95°C (10 mins), followed by 45 cycles of 95°C (15 seconds), 60°C (15 seconds) and 72°C (15 seconds). Following this, a melt curve was produced with readings taken every 1°C from 65°C to 95°C. Relative gene expression levels were normalised with the housekeeping gene 18s rRNA using the 2^−ΔΔCt^ method (Livak and Schmittgen, 2001). Normality of the data was confirmed using a Shaprio-Wilk test and equal variance confirmed using Levene’s test of homogeneity. An independent t-test was then used to compare the mean expression of each gene between the low and high risk samples and p<0.05 was considered statistically significant. All analysis was performed using SPSS (v.28.0; IBM, UK).

## Results

### RNA sequencing reveals there are differentially expressed genes between iPSC-osteoblasts derived from horses with low and high genetic risk for catastrophic fracture

An overview of the experimental design used in this study is shown in Fig. 1. The skin fibroblasts that were used in this study had previously been scored for their polygenic risk for fracture and cells with with low and high risk scores were utilised(Palomino Lago et al., 2024). Equine iPSC clones were derived in either duplicate or triplicate for each sample and were previously characterized for their ability to form embryoid bodies, express pluripotency markers, and differentiate into derivatives of endoderm, ectoderm and mesoderm (Palomino Lago et al., 2023, Bavin et al., 2015, Baird et al., 2018). All of the iPSC lines used in this study were differentiated into osteoblasts capable of producing a mineralised matrix and expressing osteoblast associated genes (Baird et al., 2018, Baird et al., 2019). RNA-sequencing was performed on 14 lines of iPSC-osteoblasts derived from six different horses (three with a low PRS (L1-3) and three with a high PRS (H1-H3) for fracture). Further information on the cells used in this study can be found in supplementary table S2.

RNA sequencing revealed that of the 29,197 mapped genes, there were 112 differentially expressed genes (DEGs) between the iPSC-derived osteoblasts from horses with a low PRS compared to those with a high PRS (Fig. 2A and Supplementary file S2). To validate the RNA sequencing results, qPCR was carried out to measure the expression of 17 genes (Fig. 2B). This demonstrated a 71% concordance between the techniques, which is in accordance with other studies (Paterson et al., 2020). Two clusters of differentially expressed genes were observed; those that were significantly more highly expressed in the osteoblasts from high risk horses (67 genes) and those that were significantly more highly expressed in the osteoblasts from low risk horses (45 genes). With the exception of samples from the low risk horse 3 (L3a and L3b), all of the samples derived from the same horse clustered together (Fig. 2 A).

**Fig. 2.**
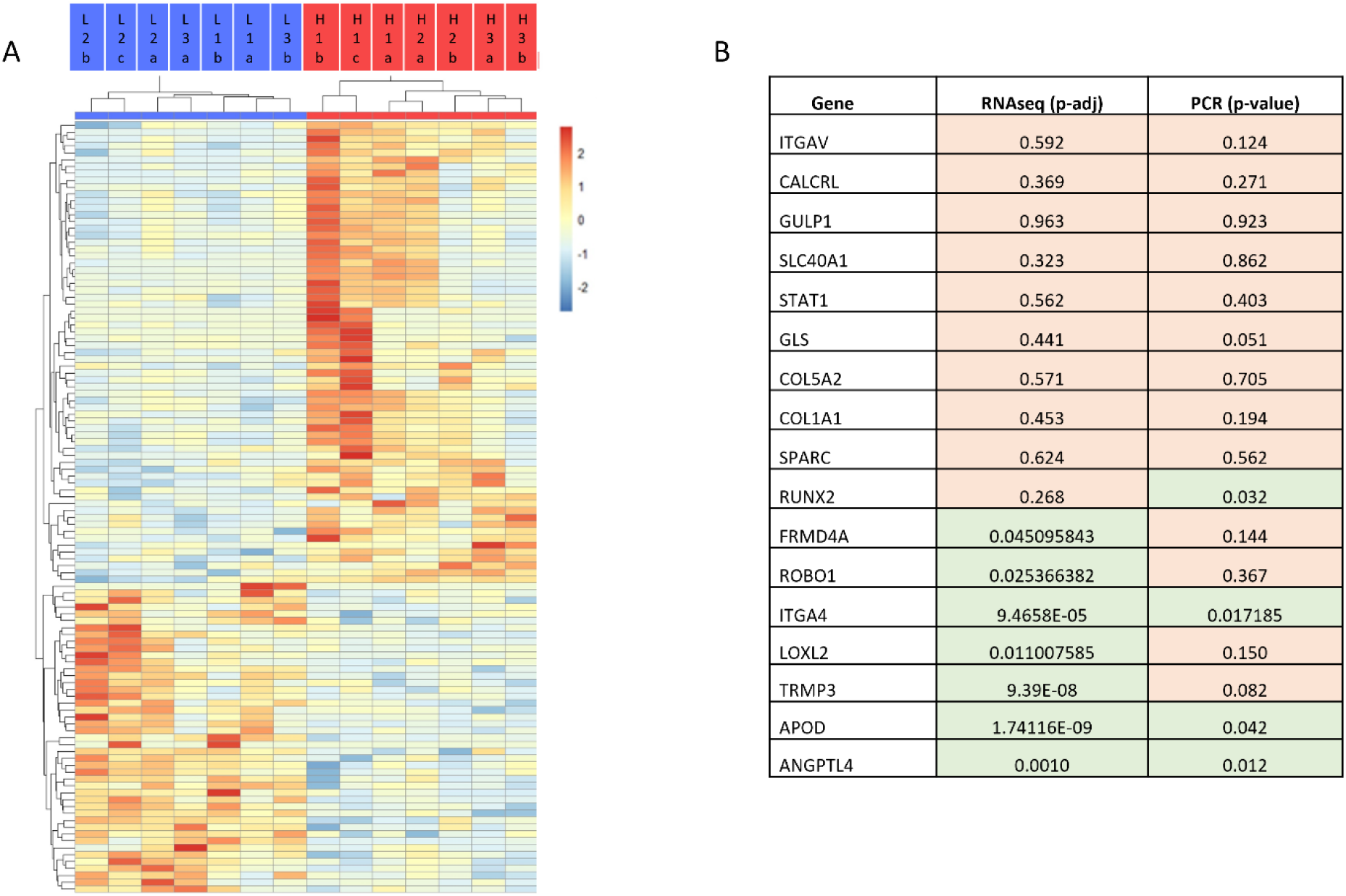
Differential gene expression in iPSC-osteoblasts derived from horses at high and low risk of fracture. A) Heatmap showing the 112 differentially expressed genes (*p*-adj<0.05 and a log2FC of ±1). The samples are shown in columns (blue = low risk samples, red = high risk samples) and genes are clustered by row (see also supplementary file S2). B) Comparison of differentially expressed genes using RNA-seq and qPCR. Genes which are expressed at significantly different levels between high and low risk samples are highlighted in green (p-value; T-test from qPCR data, or p-adj; adjusted p-value from RNAseq data <0.05), those that were not significantly different are highlighted in orange.

Of the genes that were upregulated in the low risk samples, LOC111769717 (predicted to be *PPWD1*) and LOC1000067990 (predicted to be *GBA3*) had the greatest fold changes (log2FC 23.677 and 22.306). Of the genes that were downregulated in the low risk samples, LOC111769076 (predicted to be *MUC2*), LOC111768280 (predicted to be *PRR20A*) and *PLA2G4D* had the greatest fold changes (log2FC -25.578, -23.321 and -22.848, respectively) (Fig. 3).

**Fig. 3.**
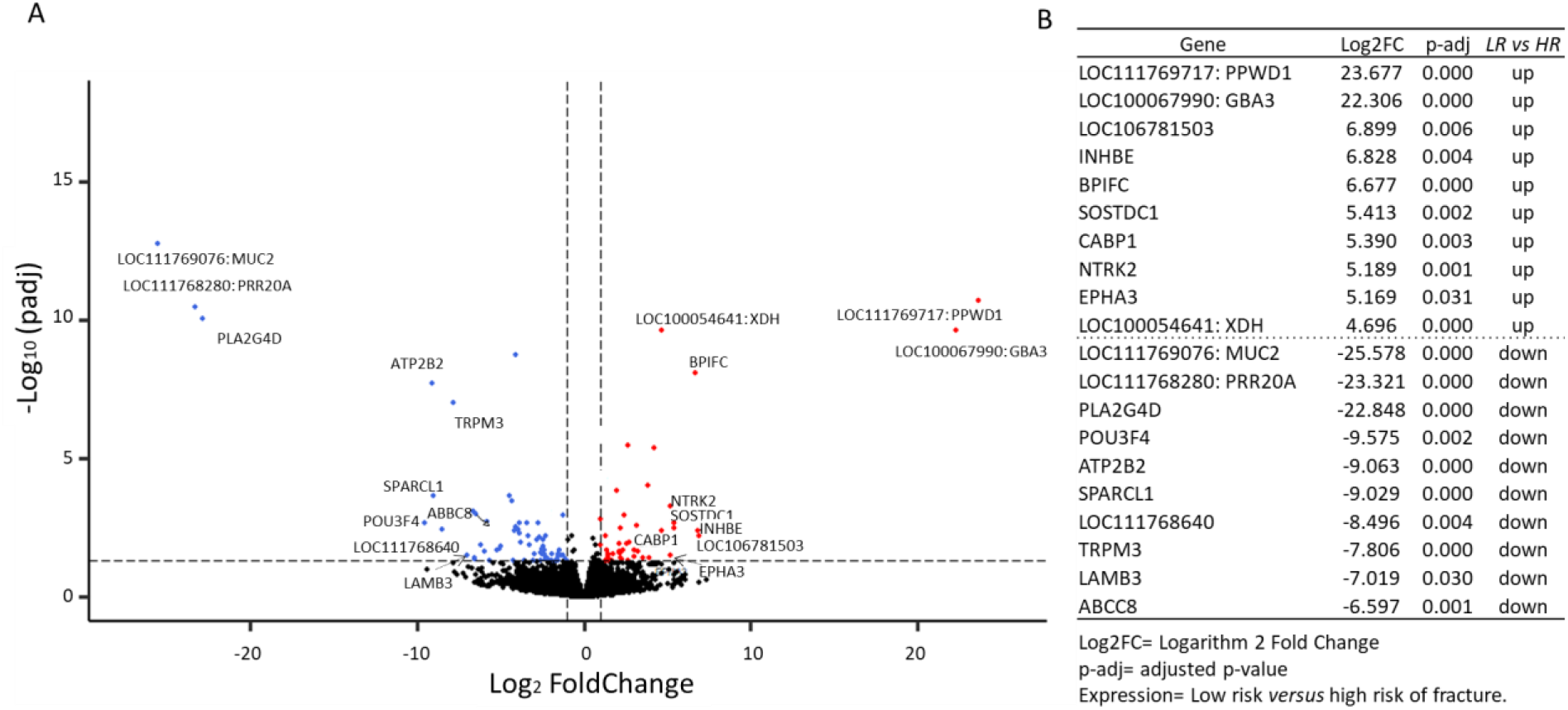
Expression of 29,197 genes in iPSC-osteoblasts from low risk compared to high risk horses. A) Volcano plot illustrating the size (log2FoldChange) and significance (*p-adj*) of the differentially expressed genes. Black dots represent genes that are not significantly different between the groups. Red dots represent genes that are significantly up-regulated in the low risk samples, blue dots represent genes that are significantly down-regulated genes in the low risk samples (*p*-adj ≤ 0.05 and log2FC of ±1). B) The top 10 up- and down-regulated differently expressed genes (DEGs) in the low risk (LR) samples compared to the high risk (HR) samples.

### A proportion of the differentially expressed genes have known roles in bone

Of the 112 DEGs, 27 are currently unannotated, 43 have published roles in bone or fracture and 42 have no known role in bone formation of fracture (Table 2). For those genes with a published role in bone or fracture further information can be found in Table 3. The DEGs were distributed across the majority of the chromosomes (Supplementary Fig. 1).

**Table 2.** The differentially expressed genes grouped by their known roles in bone, bone formation and/or fracture. C16H3orf58 = DIPK2A.

| 27 unannotated genes |  | 43 annotated genes |  |  | 42 annotated genes |  |  |
| --- | --- | --- | --- | --- | --- | --- | --- |
|  |  | Published role in bone |  |  | No published role in bone |  |  |
| LOC100054641 | LOC106781503 | ABCC5 | GPM6B | OGN | ABCC8 | ENO2 | PLPPR3 |
| LOC100067768 | LOC106781645 | ANGPTL4 | HHIP | PARVB | ABCG4 | FRMD4A | PNLIPRP3 |
| LOC100067990 | LOC106782354 | APOD | HOXD10 | POU3F4 | ABI3 | GPHA2 | PPP1R2 |
| LOC100069083 | LOC111767614 | ATP2B2 | IL15 | P2RY6 | ADGRB1 | INHBE | PTK7 |
| LOC100629234 | LOC111768280 | CHAC1 | ITGA4 | RBPMS2 | ADSSL1 | KIZ | SCNN1A |
| LOC102149222 | LOC111768640 | CHADL | LAMB3 | ROBO1 | AMPD3 | MATN4 | SCRN2 |
| LOC102149317 | LOC111769076 | CNTNAP4 | LAMC2 | ROBO2 | ASNS | MUC4 | SNAP91 |
| LOC102149614 | LOC111769201 | COBL | LGALS1 | SENP6 | BPIFC | NAP1L5 | SPARCL1 |
| LOC102149688 | LOC111769717 | COL11A2 | LOXL2 | SLC2A3 | BSN | NKAIN1 | SPRED3 |
| LOC102150273 | LOC111771940 | COL2A1 | LRRN3 | SLC38A3 | C16H3orf58 | NXPH4 | TDRKH |
| LOC102150465 | LOC111772113 | EFNB1 | MME | SMPD3 | CABP1 | PCDH9 | TMEM38A |
| LOC106781042 | LOC111774027 | EFNB2 | NR2F1 | SOSTDC1 | CERKL | PEG10 | UCP2 |
| LOC106781149 | LOC111775969 | EPHA3 | NR2F2 | SOX10 | CLHC1 | PLA2G4D | WNK4 |
| LOC106781220 |  | EYA4 | NTRK2 | TRPM3 | COTL1 | PLCH2 | ZSWIM5 |
|  |  | FBXO2 |  |  |  |  |  |

**Table 3.**
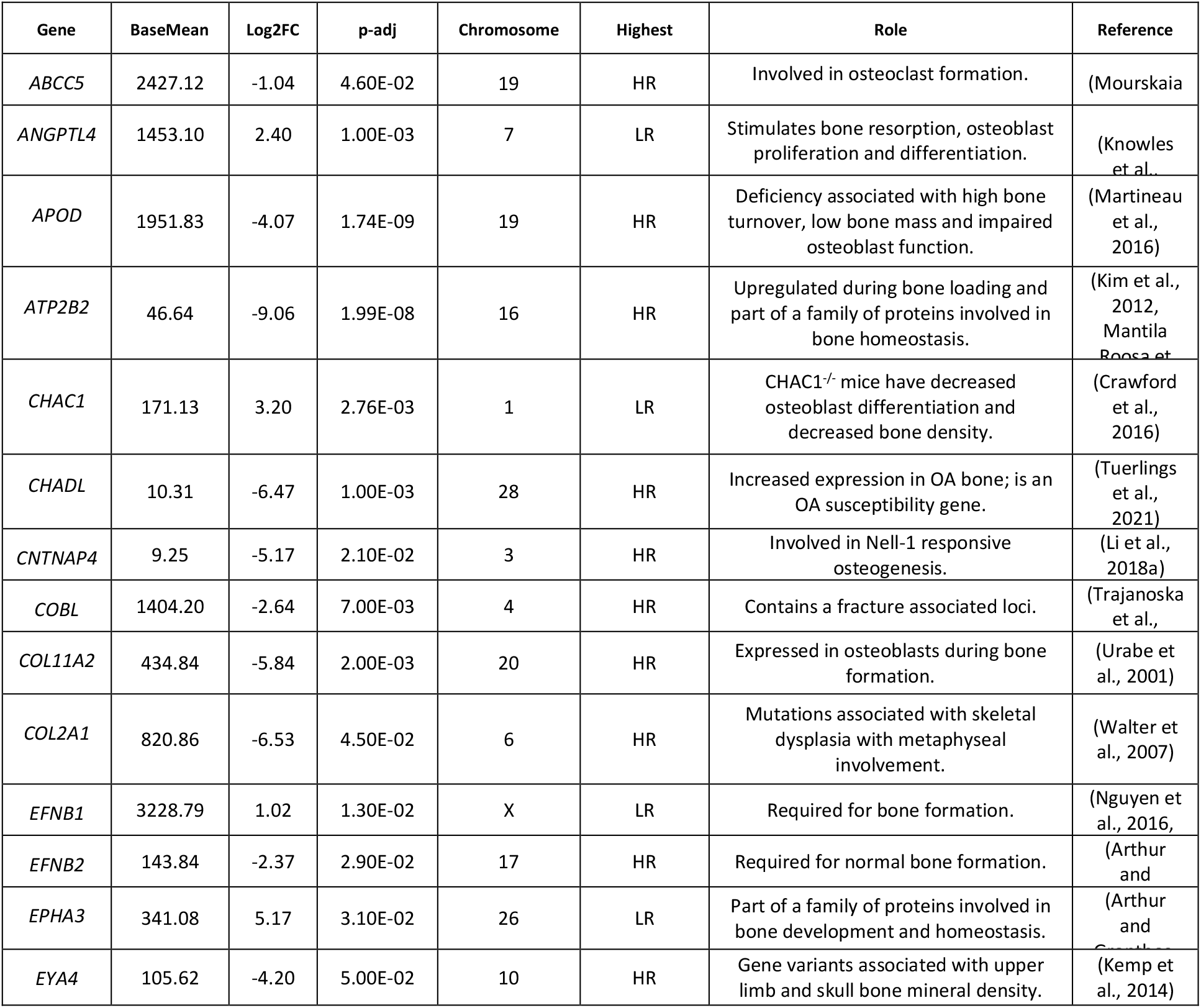
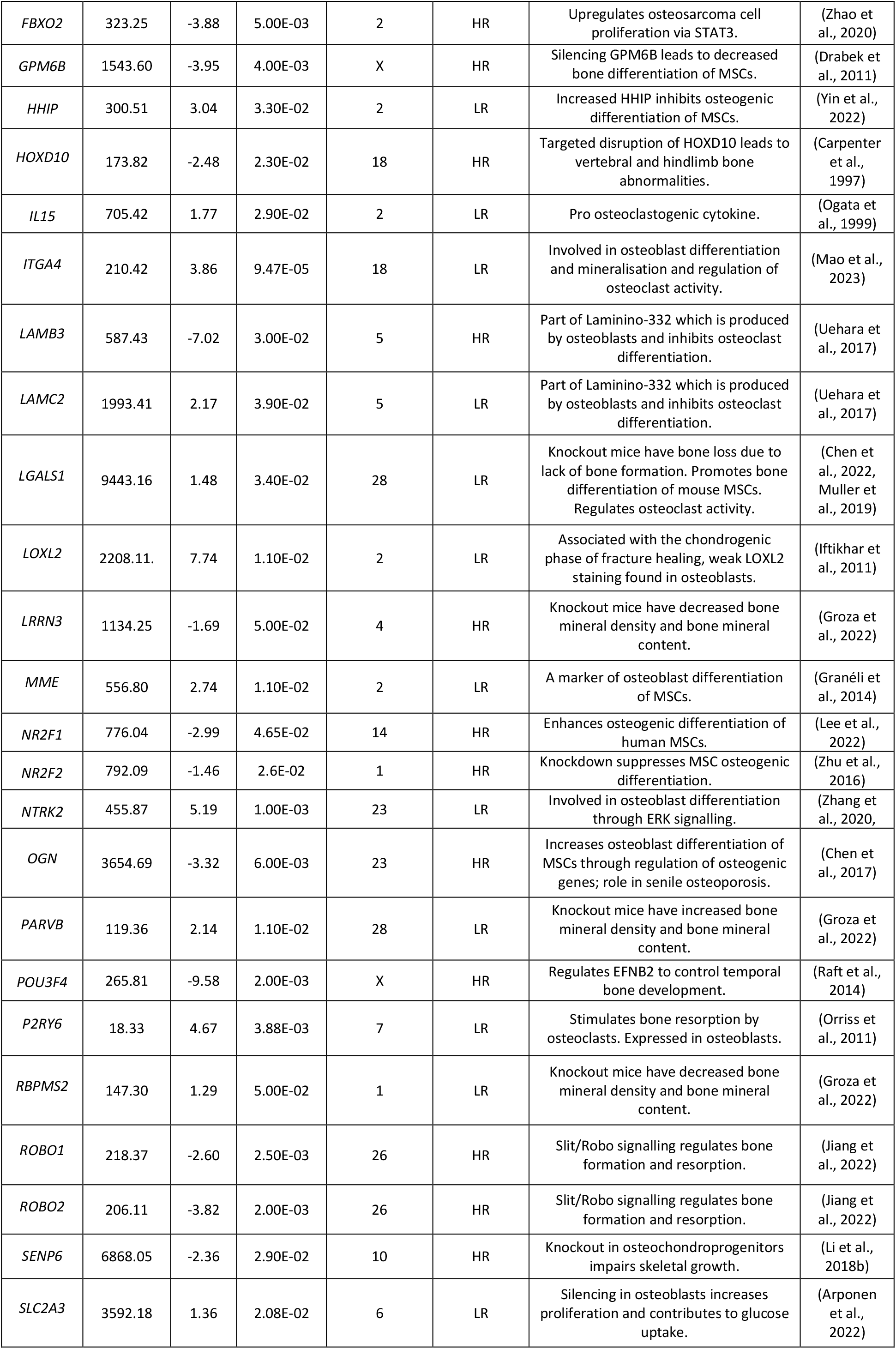
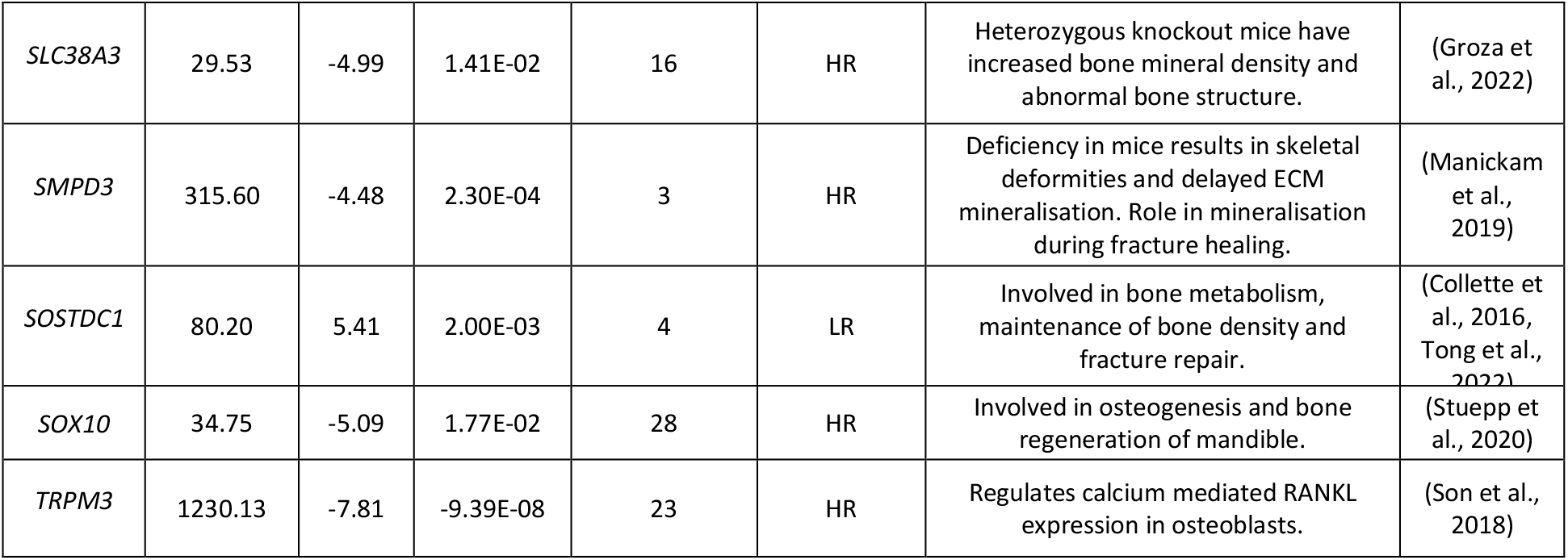
Summary of the differentially expressed genes that have a published role in bone formation or fracture. BaseMean = average of the normalised count values, dividing by size factors and taken over all samples. HR = iPSC-osteoblasts derived from high risk horses, LR = iPSC osteoblasts derived from low risk horses. OA = osteoarthritis.

However, network analysis revealed that there are many interactions between the proteins encoded by genes with known and unknown roles in bone formation and or fracture (Fig. 4).

**Fig. 4.**
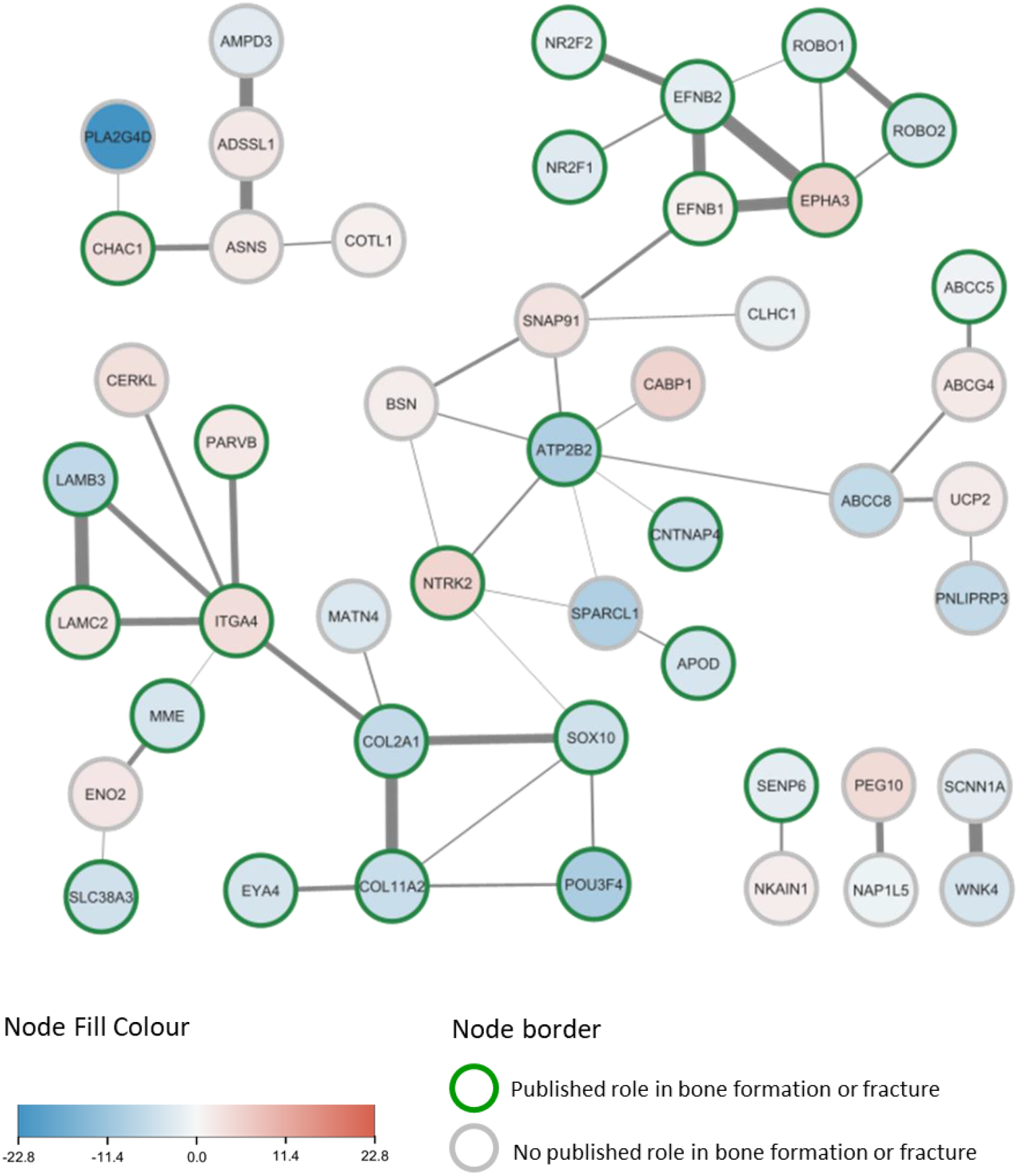
Functional network interactions of the proteins encoded by the 112 DE genes conducted with STRING. Proteins encoded by genes which are upregulated in the low risk horses are shown in shades of red, proteins which are downregulated in low risk horses are shown in blue. Proteins with a known role in bone formation/fracture have a green border, proteins with no known role in bone formation/fracture have a grey border. A thicker connecting line indicates a stronger interaction score. DEGs with zero nodes are not plotted.

### Functional enrichment of the differentially expressed genes

Gene ontology analysis of the differentially expressed genes demonstrated that they are involved in numerous biological processes including adhesion, development, morphogenesis, differentiation and extracellular matrix organisation. Many of the differentially expressed gene products were also located in the extracellular matrix (Fig. 5).

**Fig. 5.**
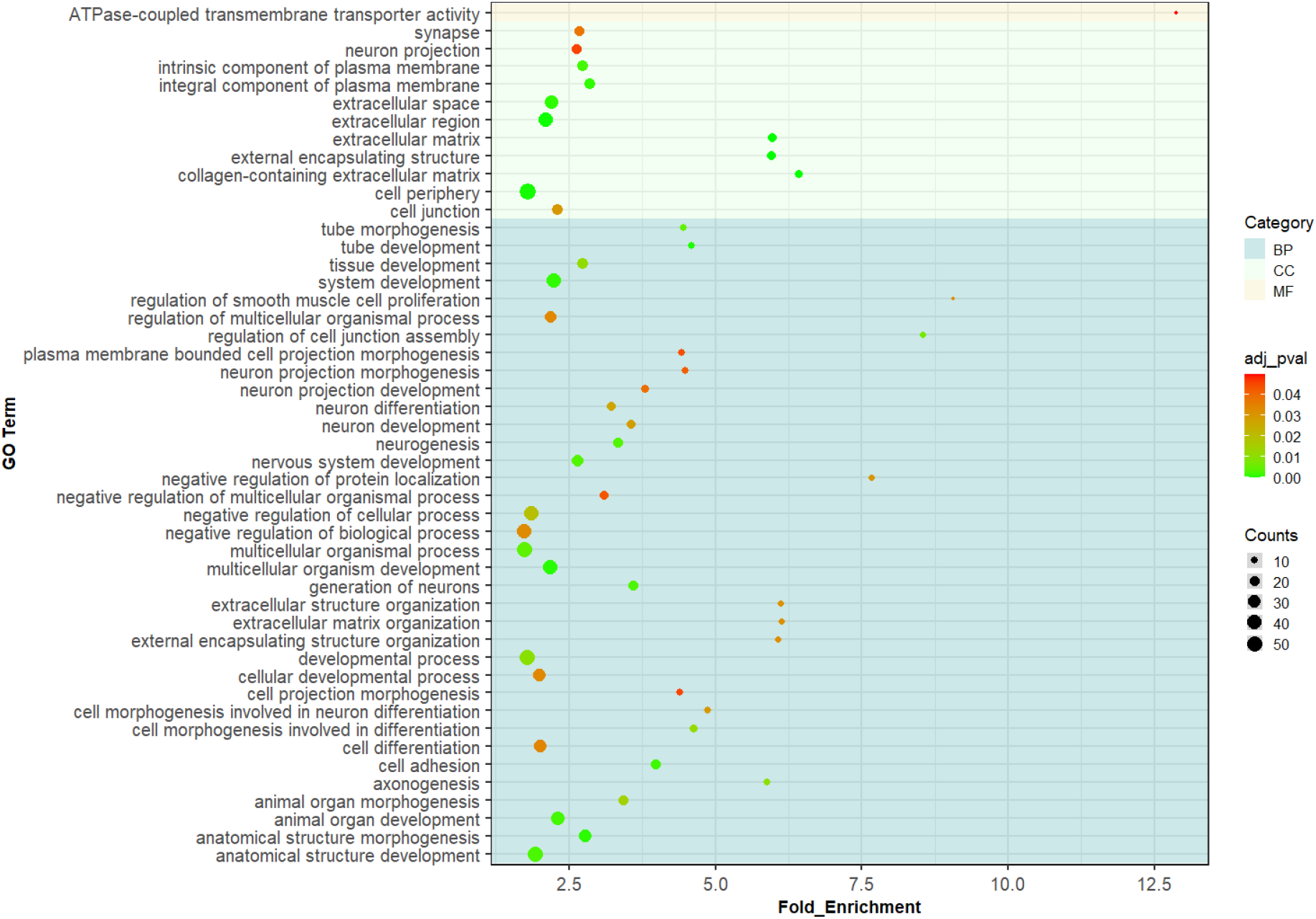
Gene ontology (GO) terms significantly over-represented by the differentially expressed genes. Background colour corresponds with Biological Processes (BP – dark blue), Cellular Component (CC – light blue) and Molecular Function (MF – light orange). The colour of the dot shows the significance of the GO terms (colour scale bar indicating the range from 0 (green) to 0.04 (red) adjusted p-value). The size of the dot represents the number of DEGs in that specific GO term.

Pathway analysis was also performed to identify pathways that are over-represented by the differentially expressed genes (Fig. 6). This revealed a range of affected pathways including the ERK pathway, degradation of the extracellular matrix and collagen chain trimerization.

**Fig. 6.**
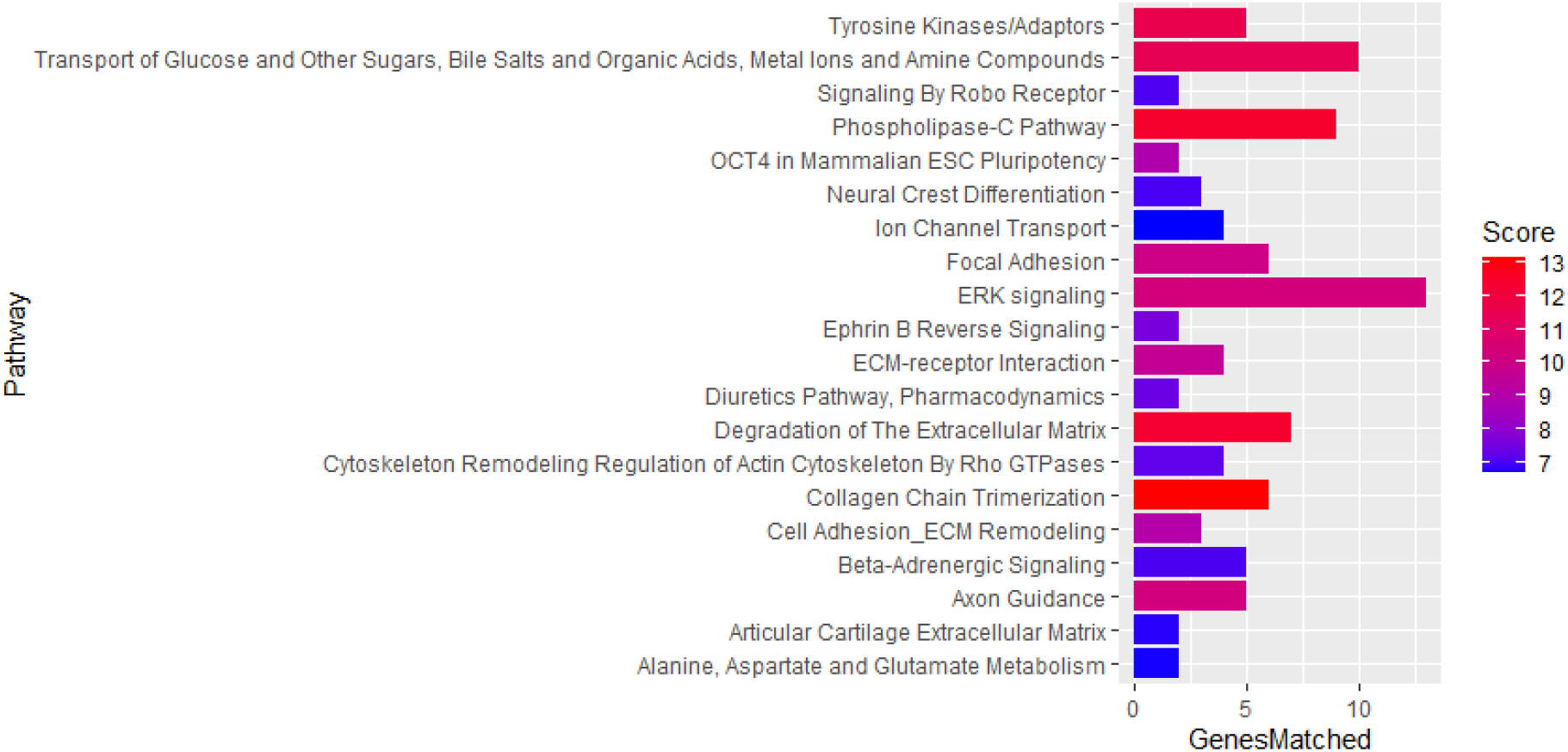
Pathways that are significantly over-represented by the DEGs. Score is a transformation of the binomial distribution p-value and the score range is divided into three quality levels, according to the p-value they are derived from (after correction for multiple comparisons), represented from lower (blue) to higher (red) score. GenesMatched = number of genes over-represented in that specific pathway.

### Gene Set Enrichment Analysis GSEA

As over-representation analysis was performed on the DEGs only, it excludes genes that did not pass the original cut-off (p-adj <0.05, log2FC ±1) but that could have a relevant function in a complex disease. Gene Set Enrichment Analysis (GSEA) associates a disease phenotype (high risk of fracture) to a group of genes/proteins (Subramanian et al., 2005, Mootha et al., 2003). GSEA demonstrated that 6 GO-based gene sets, 6 KEGG (Kyoto Encylopedia of Gene and Genomes)-based gene sets, 11 WP (WikiPathways)-based gene sets and 4 REACTOME-based gene sets, were significantly enriched (Table 4) (FDR<0.05). Many of the enriched pathways and processes were associated with glycolysis, and the associated genes had a higher expression in the iPSC-osteoblasts from horses with low fracture polygenic risk scores.

**Table 4.**
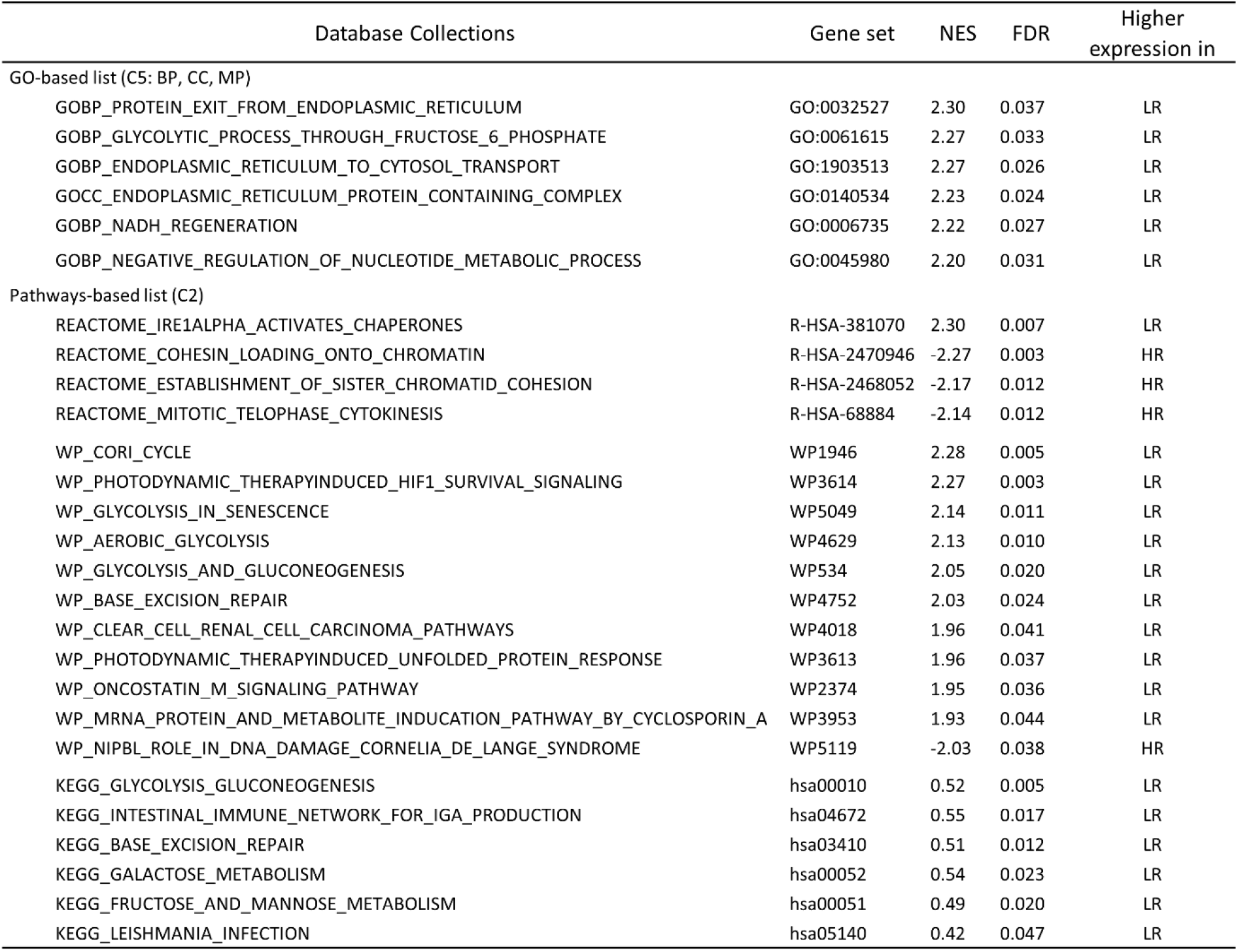
Gene set enrichment analysis (GSEA) showing the significantly enriched gene ontology terms and pathways. BP=biological process, CC=cellular component, MP=molecular process. WP=WikiPathways, KEGG=Kytoto Encyclopaedia of Genes and Genomes. NES=normalised enrichment score. FDR=False discovery rate.

## Discussion

In this study we identified 112 genes that were differentially expressed between iPSC-osteoblasts derived from horses at high and low genetic risk of fracture. Of these, 27 are unannotated, 43 have published roles in bone formation or fracture and 42 have no known role in bone formation of fracture. However, of these 42 genes, all but five of them have been reported to be expressed in equine bone tissue in publicly available datasets (Kuemmerle et al., 2016, Kemper et al., 2019). Furthermore, STRING analyses demonstrated that 18 of the 42 genes with no known role in bone encode proteins which interact (directly or indirectly) with proteins that have known roles in bone. A further four of the unknown genes interact with each other. This suggests that at least a proportion of these genes are likely to have roles in bone which have not yet been discovered and should be followed up in future work.

Of the top 10 genes with the largest DE that were upregulated in the LR samples four of them are not yet annotated in the horse genome. However, LOC111769717 is predicted to be *PPWD1*, whose expression is associated with post-menopausal osteoporosis (Qian et al., 2019). Two of the other non-annotated transcripts also have predicted genes (*GBA3* and *XDH*), however a role in bone has not been reported. Three of the other top 10 genes have some previously published links with bone. For example, *SOSTDC1* has been shown to be involved in bone metabolism, the maintenance of bone density and fracture repair (Tong et al., 2022, Collette et al., 2016). *NTRK2* is involved in osteoblast differentiation of MSCs (Zhang et al., 2020) and binding of NTRK2 to its receptor results in the phosphorylation of ERK1/2 which drives the expression of the bone-promoting transcription factors *RUNX2* and *SP7* (Osterix) (Liu et al., 2018). *EPHA3* is a member of the ephrin receptor family and ephrin signalling is involved in bone development and homeostasis (Arthur and Gronthos, 2021).

Two of the genes, *INHBE* (a member of the TGF-β superfamily) and *BPIFC* (predicted to enable lipid binding) have no known role in bone formation or fracture, do not appear in the STRING analysis as associating at the protein level with the other differentially expressed encoded proteins, and they are not in the gene sets of the enriched GO terms or pathway analysis. Therefore, it is not clear what their biological relevance might be. However, the expression of both genes has been detected in horse bone tissue, and/or or cultured horse cells (Kuemmerle et al., 2016). *CABP1* is also in the top 10 and although it has no published role in bone, it appears in the main interaction node of the STRING analysis, is a calcium binding protein and is expressed in horse bone tissue (Kuemmerle et al., 2016, Kemper et al., 2019).

Of the top ten genes that were downregulated in the LR samples, three were not annotated and, where available, their predicted genes have no links to bone or fracture. Three of the genes, *PLA2G4D*, *SPARCL* and *ABCC8* have no known role in bone formation or fracture. All three genes are expressed in horse bone tissue, and/or or cultured horse cells (Kuemmerle et al., 2016). ABCC8 functions as a modulator of ATP-sensitive potassium channels and insulin release and it is in the enriched gene set for the overrepresented pathway transport of glucose and other sugars, bile salts and organic acids, metal ions and amine compounds. Furthermore, it interacts with ATB2BP2, which is also significantly downregulated in the LR samples. SPACRL also interacts with ATB2BP2. SPARCL, which is predicted to enable calcium ion, collagen and extracellular matrix binding activity, is a paralog of SPARC, which plays a critical role in bone remodelling (Rosset and Bradshaw, 2016). However, the role of SPARCL in bone is so far unknown. PLA2G4D also appears in the STRING network and interacts with CHAC1. Knock-out of *CHAC1* in mice leads to decreased osteoblast differentiation and bone density (Crawford et al., 2016). PLA2G4D is part of the phospholipase A2 family, and it is present in the enriched gene set for the overrepresented pathway phospholipase-C pathway.

The remaining five genes of the top ten that were downregulated in the LR samples include *TRPM3* which is involved in bone remodelling through the regulation of RANKL expression in osteoblasts (Son et al., 2018). Similarly, *LAMB3* is also involved in bone remodelling. LAMB3 forms part of laminin-332 which is produced by osteoblasts and supresses osteoclast differentiation (Uehara et al., 2017). *ATP2B2* is involved in maintaining calcium homeostasis. It is upregulated during bone loading (Mantila Roosa et al., 2011), expressed by osteoblasts (Francis et al., 2002) and is part of a family of proteins which are required for regulating bone mass (Kim et al., 2012). *POU3F4* regulates *EFNB2* to control temporal bone development (Raft et al., 2014). Interestingly, *EFNB2* itself was also significantly downregulated in the LR samples and the ephrin B reverse signalling pathway was significantly over-represented by the differentially expressed genes.

Therefore, the known functions of the top 10 up and down regulated genes, suggest that the regulation of bone remodelling and calcium signalling may be altered in bone cells from horses at high risk of fracture.

GO analysis of all the differently expressed genes revealed that the molecular function “ATPase-coupled transmembrane transporter activity” was over-represented. ATPase activity is crucial for regulating bone formation and resorption (Francis et al., 2002). GO analysis further revealed that the extracellular matrix/region was most often over-represented by the differently expressed genes. For the biological processes, in addition to many processes involved in differentiation and development, there were a number of processes related with the extracellular matrix (ECM). Together this may suggest that the differentially expressed genes are ultimately affecting bone homeostasis and ECM components. It is noted that many neuronal GO processes are also over-represented by the DEGs. The cross-talk between brain and bone health is becoming increasingly apparent (Otto et al., 2020) and it is possible that many of the DEGs have to date been better studied in neuronal cell types rather than osteogenic cells, influencing the GO processes they are known to be involved in. For example, Slit/Robo signalling was initially found to be essential during nerve development, before its role in regulating bone formation was discovered (Jiang et al., 2022). Therefore, *ROBO1*/*ROBO2* appear in many of the gene lists associated with neuronal biological processes and pathways.

Pathway analysis revealed multiple pathways involved in bone. For example, the ERK signalling pathway had the greatest number of DEGs and this pathway is involved in fracture repair (Chen and Luan, 2019). Collagen chain trimerization was one of the most significantly over-represented pathways and this is affected in numerous bone diseases (Bourhis et al., 2012). Similar to the GO analysis, pathways relating to the degradation and remodelling of the ECM were also over-represented. GSEA revealed biological processes and numerous pathways involved in glycolysis. Human patients with diabetes are at higher risk of bone fractures (Valderrábano and Linares, 2018) and altered glucose metabolism may be important in maintaining bone homeostasis (Karner and Long, 2018).

This study had a number of limitations. Osteoblasts were only derived from a small number of horses. This may have resulted in a lack of power to detect additional significant DEGs. However, due to the samples we had available (Palomino Lago et al., 2024), we were not able to increase the sample size whilst selecting samples that were at the most extreme ends of the polygenic risk score. Furthermore, in our previous study (Palomino Lago et al., 2024), we demonstrated that fibroblast cells differentiated directly into osteoblast-like cells, had a significant difference in the expression of *COL3A1* and *STAT1* between samples isolated from high and low risk horses (Palomino Lago et al., 2024). In this study, the reference transcriptome used did not have *COL3A1* annotated, and while we saw the same trend for *STAT1*, the difference was not significant (Supplementary file 2). This may reflect the different starting populations of cells and the fact that the osteoblast populations we derived in both studies are likely to be heterogeneous. The *COL3A1* and *STAT1* genes lie within a fracture associated region on ECA18 (Blott et al., 2014, Tozaki et al., 2020). In this study, we found only three of the differentially expressed genes to be located on ECA18 (*HOXD10*, *ITGA4*, *CERKL*). However, none of them lie within the associated region. It is not clear if our study lacked the power to detect smaller differences in the expression of any of these genes from this region, or if DNA variants within the region are regulating more distant genes. Finally, in our model, we directed the iPSCs to differentiate into osteoblasts. However, it is likely that other cell types (e.g. osteoclasts) also contribute to fracture risk. Human iPSCs have successfully been differentiated into osteoclasts (Chen, 2020), but to date this not yet been reported for equine iPSCs and it would be of benefit to measure gene expression profiles in other cell types in the future. Similarly, we did not have access to bone tissue from fracture cases and controls and so were unable to confirm if these differences in expression also occur *in vivo* at either the gene or protein level.

In conclusion, we have demonstrated that iPSC-osteoblasts derived from horses with high and low polygenic risk scores for catastrophic fracture, have many differently expressed genes that are overrepresented in various pathways and processes that have relevance to bone homeostasis and fracture. A deeper understanding of the consequences of mis-regulation of these genes and the identification of the DNA variants which underpin their differential expression may reveal more about the molecular mechanisms which are involved in equine bone health and fracture risk.

## Acknowledgements

This study was kindly funded by the Horserace Betting Levy Board (vet/prj/792). The Alborada Trust fund A.K.C.R.

## Data availability

The RNA sequencing datasets are available in the National Centre for Biotechnology Information Gene Expression Omnibus repository (NCBI, GEO www.ncbi.nlm.nih.gov/geo) under accession number GSE255417. The differentially expressed genes and normalised counts data are included in Supplementary File S2.

## Supporting information captions

**Supplementary Table S1.** Forward and reverse primer sequences used for RT-qPCR analysis.

**Supplementary Table S2**. Summary of the iPSCs used in this study. H1-3 are samples from three high risk (HR) horses. L1-3 are samples from three low risk (LR) horses. iPSCs were derived in duplicate or triplicate from each horse (a, b, c). The polygenic risk scores (PRS) for fracture of the donor horses are provided (Palomino Lago et al., 2024) along with the passage at which they were differentiated into osteoblasts for use in RNA sequencing.

**Supplementary Fig. S1.** The distribution of the annotated differentially expressed genes (DEGs) across the equine chromosomes. Green bars represent those genes with a published role in bone, grey bars represent those genes with no published link to bone.

**Supplementary File S2.** Gene level summaries providing the normalised count data and differential expression levels.

