## Supplementary data for "Identification of a global gene expression signature associated with the genetic risk of catastrophic fracture in iPSC-derived osteoblasts from Thoroughbred horses"

**Supplementary Figure 1**

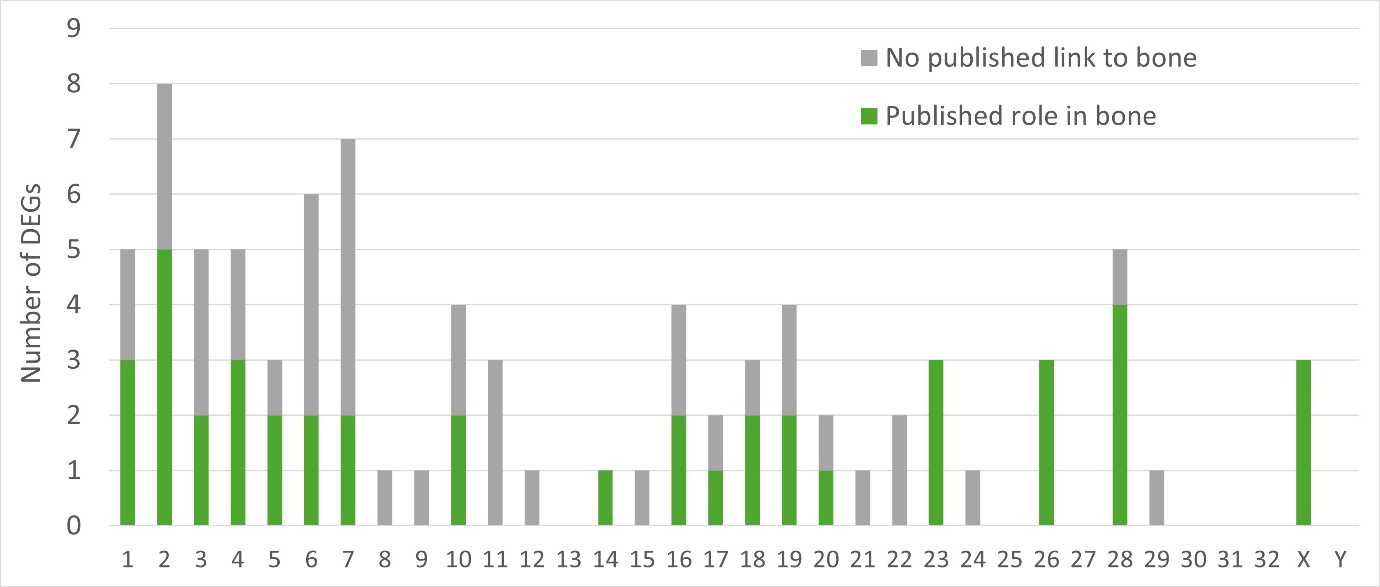

**Supplementary Figure 1.** The distribution of the annotated differentially expressed genes (DEGs) across the equine chromosomes. Green bars represent those genes with a published role in bone, grey bars represent those genes with no published link to bone.

**Supplementary Table S1**

| **Gene** | **Protein Name** | **Forward Primer (5’-3’)** | **Reverse Primer (5’-3’)** |
| --- | --- | --- | --- |
| *18S rRNA* | 18s ribosomal RNA | CCCAGTGAGAATGCCCTCTA | TGGCTGAGCAAGGTGTTATG |
| *ITGAV* | Integrin Subunit Alpha V | GATGCTGATGGACAGGGATT | AAACTACCAGGACCACCAAGAA |
| *CALCRL* | Calcitonin Receptor Like Receptor | TTTGTGTTTCTCTTGCCTTTTT | TCTTCTGATTCTGCTGTGACAA |
| *GULP1* | GULP PTB Domain Containing Engulfment Adaptor 1 | GCCCAAAGGAACAGAAGTTG | GGAATTTTCTGGCCTTCAGA |
| *SLC40A1* | Solute Carrier Family 40 Member 1 | GCCAACTACCTGACCTCTGC | AAAGTGCCACATCCGATCTC |
| *STAT1* | Signal Transducer And Activator Of Transcription 1 | TCTGCAGCTGTCTGAAGGAG | TGAATATTCCCCGACTGAGC |
| *GLS* | Glutaminase | CCTAGCTTGGAAGATTTGCTG | CAGACGTTCGCAATCCTGTA |
| *COL5A2* | Collagen Type V Alpha 2 Chain | AGGAGAGAGAGGCCCAAAAG | CTCCATCAATTCCCTGAGGA |
| *COL1A1* | Collagen Type I Alpha 1 Chain | TGCGAAGACACCAAGAACTG | GACTCCTGTGGTTTGGTCGT |
| *SPARC* | Secreted Protein Acidic and Cysteine Rich | TGGCGAGTTTGAGAAGGTGT | TTTGCAAGGCCCGATGTAGT |
| *RUNX2* | RUNX Family Transcription Factor 2 | CCAAGTGGCAAGGTTCAACG | AACTCTTGCCTCGTCCACTC |
| *FRMD4A* | FERM Domain Containing 4A | AGGGCCGTCGATGTCAAG | AGCCAGTTTAAGTGTCCCGT |
| *ROBO1* | Roundabout Guidance Receptor 1 | AACACCAGCCAGGACATCTG | TAGACGGGAGTTTTGGCACC |
| *ITGA4* | Integrin Subunit Alpha 4 | GAGGAGGGGAGAGTGTTCGT | TTGCAGCGTATTTGTCGCTTC |
| *LOXL2* | Lysyl Oxidase Like 2 | AAGCCTACAAGCCAGAGCAAC | CACCAAGTCCCACTTATCGTCA |
| *TRPM3* | Transient Receptor Potential Cation Channel Subfamily M Member 3 | TGCCTGCCGTTTTTCTCTCT | AACGCCTTGACCGATTCCAA |
| *APOD* | Apolipoprotein D | GCATGTGGAAACTGCCTTCAT | CATCACCATCCTGGCGTTGG |
| *ANGPTL4* | Angiopoietin Like 4 | GAGAAGCAGCGCCTGAGAAT | CCTCCTCCCGGCAGACTTA |

**Supplementary Table S1.** Forward and reverse primer sequences used for RT-qPCR analysis

**Supplementary Table 2**

| **Sample** | **Group** | **PRS** | **Cell Passage** |
| --- | --- | --- | --- |
| **H1a** | **HR** | **1.48E-06** | **13** |
| **H1b** | **HR** | **1.48E-06** | **13** |
| **H1c** | **HR** | **1.48E-06** | **14** |
| **H2a** | **HR** | **1.15E-06** | **7** |
| **H2b** | **HR** | **1.15E-06** | **8** |
| **H3a** | **HR** | **1.05E-06** | **9** |
| **H3b** | **HR** | **1.05E-06** | **7** |
| **L1a** | **LR** | **-2.72E-06** | **10** |
| **L1b** | **LR** | **-2.72E-06** | **9** |
| **L2a** | **LR** | **-3.73E-06** | **14** |
| **L2b** | **LR** | **-3.73E-06** | **16** |
| **L2c** | **LR** | **-3.73E-06** | **10** |
| **L3a** | **LR** | **-2.57E-06** | **9** |
| **L3b** | **LR** | **-2.57E-06** | **10** |

**Supplementary Table S2.** Summary of the iPSCs used in this study. H1-3 are samples from three high risk (HR) horses. L1-3 are samples from three low risk (LR) horses. iPSCs were derived in duplicate or triplicate from each horse (a,b,c). The polygenic risk score (PRS) for fracture of the donor horses are provided (18) along with the passage at which they were differentiated into osteoblasts for use in RNA sequencing.
